# Brain FACTs: A same-section workflow for Fiber Architecture, Cytoarchitecture and cell Types in large-scale brain sections

**DOI:** 10.64898/2026.10.13.758849

**Authors:** Roxana N. Kooijmans, Markus Axer, David Gräßel, Katrin Amunts

## Abstract

Understanding the human brain’s multiscale organization requires linking cells, connections and architecture, yet conventional methods are incompatible within a single specimen, and inter-individual variability confounds comparisons. Multiple-label immunohistochemistry has been applied in interleaved sections but remains challenging in single large sections and has never been combined with 3D polarized light imaging (3D-PLI) of the fiber architecture. Here we introduce a sequential multimodal workflow that integrates 3D-PLI, multiplex chromogenic immunohistochemistry, and Nissl staining on the same large human brain section, while fully preserving each technique’s quality and specificity. The workflow was validated on sections from vervet monkey and Wistar rat. The resulting paired high-resolution images enable direct micron-scale mapping of fiber tracts relative to molecularly defined cell types, yielding previously inaccessible insights including molecular identification of white-matter components, refined cortical layering and cytoarchitectural disambiguation. This approach bridges spatial scales, clarifies relationships between modalities, and reveals organizational similarities and differences across species.

---

Investigating the human brain, from basic research to therapeutic targeting, requires detailed insight into its microarchitecture. This knowledge remains incomplete, largely due to methodological limitations. While new molecular and genetic markers continue to emerge from animal models^1^, they only widen rather than close the gap, since direct generalization to human is limited by divergent evolution^2^, brain size, and organizational complexity. Although many cell types exhibit homology across species^3^ data obtained in mice and other models cannot be directly transferred to the human brain. Generating high-resolution maps of the human brain for novel molecular targets, ideally embedded in standardized parcellations, is therefore essential to strengthen the translational relevance of animal findings.

Significant advances have been made in spatial transcriptomics, yet protein expression maps are still largely missing. Protein levels are more directly linked to function, with reported prediction value from transcription ranging between 15 and 85%^4^. Several challenges have slowed progress in this area. Experimental data across different species remain fragmented, partly because methodological differences prevent direct comparison. Extensive cellular and connectivity datasets exist for mouse^5^, including full-brain coverage, whereas corresponding information for the human brain is still incomplete^6^. The human brain is more than 3500 times heavier and contains more than 1200 times as many neurons than a mouse brain^7^, posing substantial technical challenges. The BigBrain^8^ data set a new standard for uniform, high resolution anatomical model of the human brain at near-cellular resolution, but it lacks cell-type specific information and connectivity information due to incompatible methods.

Silver staining for cell bodies^9^, classical Nissl^25^ and myelin stains^10^ provide robust measures of overall cyto- and myeloarchitecture, yet they aggregate signals from mixed cell and fiber populations. Classical myelin staining, moreover, obscures fiber direction due to the densely packed fibers within a section, a major limitation for studying connectivity at the cellular scale. This problem has been addressed by 3D Polarized Light Imaging (3D-PLI)^11^, which exploits the birefringence properties of myelin to map the three-dimensional course of axons throughout the brain. 3D-PLI has been successfully applied in many species, including rat^12^, vervet monkey^13^, and human^14^, an important prerequisite for inter-species comparisons.

At the cellular level, the prevalence and spatial distribution of cell types differ across species and brain regions, as shown by differences in epigenetic^15^, transcriptomic^16^, proteomic^17^, morphological^18^, connectomic,^19^ and, electrophysiological studies^20^ . Cell censuses^21^ provide quantitative baselines but fall short in topographic mapping and relating cell types to connectivity or fine-grained parcellation.

Classical^22^ and recent^23^ studies exploring different immunomarkers used interleave sections^24^ with small intervals to analyze the relationships of different cell types. A few recent studies have reported multiplex immunohistochemistry in human brain sections: Cyclic multiplex fluorescent immunohistochemistry on human postmortem brains has been used for a quantitative characterization of the morphology of astrocytes and microglia in sections of the temporal pole of patients with Alzheimer and controls^25^. Another study showed triple enzymatic immunohistochemistry for parvalbumin, calbindin, and calretinin interneurons from samples of different cortical areas of the human brain^26^.

Although multimodality is widely recognized as essential for a comprehensive understanding of brain organization^27^, registering data acquired with different methods remains challenging because of inter- and intra-species structural variability^28^, particularly the complex folding of the human cerebral cortex^28^. Spatial reference spaces help integrate modalities^29^, yet intersubject differences in brain macroscopy limit precision at the cellular level.

Here we introduce FACTs, a mapping approach that overcomes some of these limitations by integrating micro-scale fiber architecture, cytoarchitecture and molecularly identified cell types, in one and the same large-scale brain sections of human, rat and vervet brains. We adapted (immuno)histological and imaging protocols into a sequential workflow that combines label-free 3D-PLI^14^, multiplex chromogenic immunohistochemistry, and Nissl staining^30^ on identical large-scale sections while preserving the quality and specificity of each modality. We developed solutions for image fusion, as well as data handling of images in the 10 GB range per modality plane (see Online Methods). The resulting co-registered data sets enable, for the first time, direct topographic analysis of the relationships between fibers, molecularly defined cells, and cytoarchitectures across the human brain. The method has also been applied in vervet monkey and rat brain tissue.

## Results

We developed an integrated histology and imaging workflow: FACTs, standing for Fibers, Architecture and Cell Types (Fig. 1) that enables micron-resolution, same-section acquisition of fiber orientation, cytoarchitecture and molecularly defined cell types across large-scale brain sections (Fig. 2). The method was applied to full human brain sections and successfully replicated in vervet monkey and rat brain sections (Supplementary Table 1). The workflow is applicable to formaldehyde-fixed tissue, whether fixed in formalin or in paraformaldehyde (PFA), making it accessible for regularly preserved human post-mortem tissue, as well as experimental animal tissue. Fixation is followed by washing in running tap water for 1 week for the full human brain, and moving the tissue to phosphate buffered saline. After that, T2* weighted MR images (Philips Ingenia 3.0T CX) are acquired to correct for tissue distortions and to facilitate reconstruction of the full brain volume^8^, for digital atlas registration^31^. The frozen hemispheres or brains are sectioned on a Leica Biosystems CM3600 freezing microtome. Polarized light imaging measurements on a Taorad LMP-1 microscope have been applied^32^, followed by tissue re-floating, chromogenic immunohistochemistry, and Nissl staining, with brightfield imaging on a Huron TissueScope™120 Slide Scanner. Data have been stored at the Jülich Supercomputing Centre. Multiplex chromogenic immunohistochemistry for calcium binding proteins (parvalbumin (PV), calbindin (CB), calretinin (CR)), and Nissl staining on identical sections have been performed on one and the same section (Fig. 2). This approach embeds axonal architectures and molecularly labeled cells directly into the cytoarchitectonic context of the same section, spanning multiple spatial scales - from the section overview, through cytoarchitectonic areas, e.g., area PG in the posterior inferior parietal lobule^33^ as illustrated in Fig. 2.

**Fig. 1.**
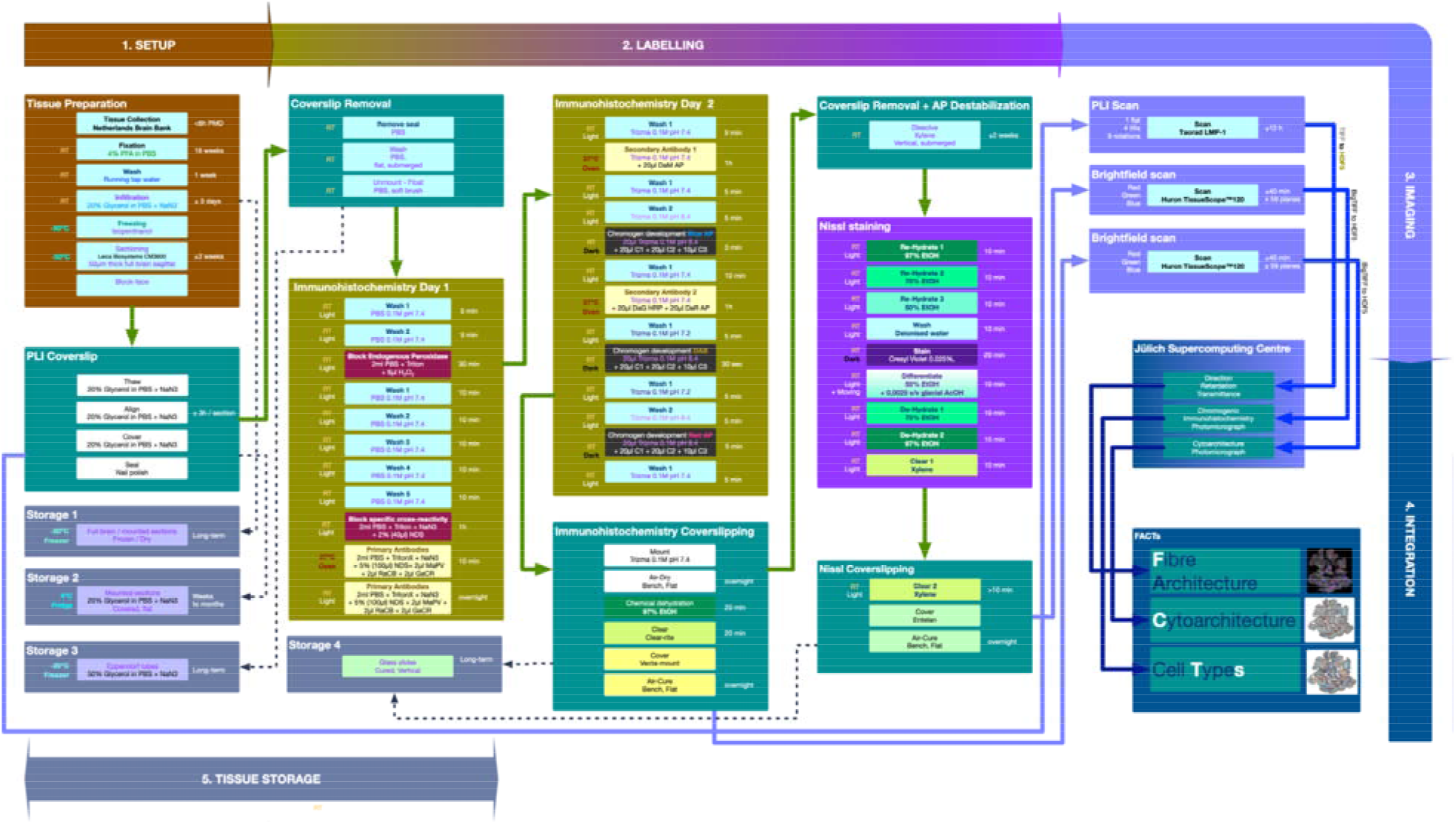
Overview of the FACTs workflow, showing the principal processing stages and the corresponding state of the tissue at each stage. (1) General tissue preparation: sectioning of the whole brain on a Leica Biosystems CM3600 freezing sliding microtome. (2) Labelling: re-floating of the sections following 3D-PLI, and sequential multiplex chromogenic immunohistochemistry and Cresyl violet Nissl staining. (3) Imaging: label-free 3D polarized light imaging (3D-PLI) of the unstained sections on a Taorad LMP-1 microscope, and brightfield imaging of the labeled sections on a Huron TissueScope™120 slide scanner. (4) Data integration: storage and processing of the same-section image sets at the Jülich Supercomputing Centre. (5) Storage: a holding stage, accessible from every other stage, that accommodates their differing durations and throughput.

**Fig. 2.**
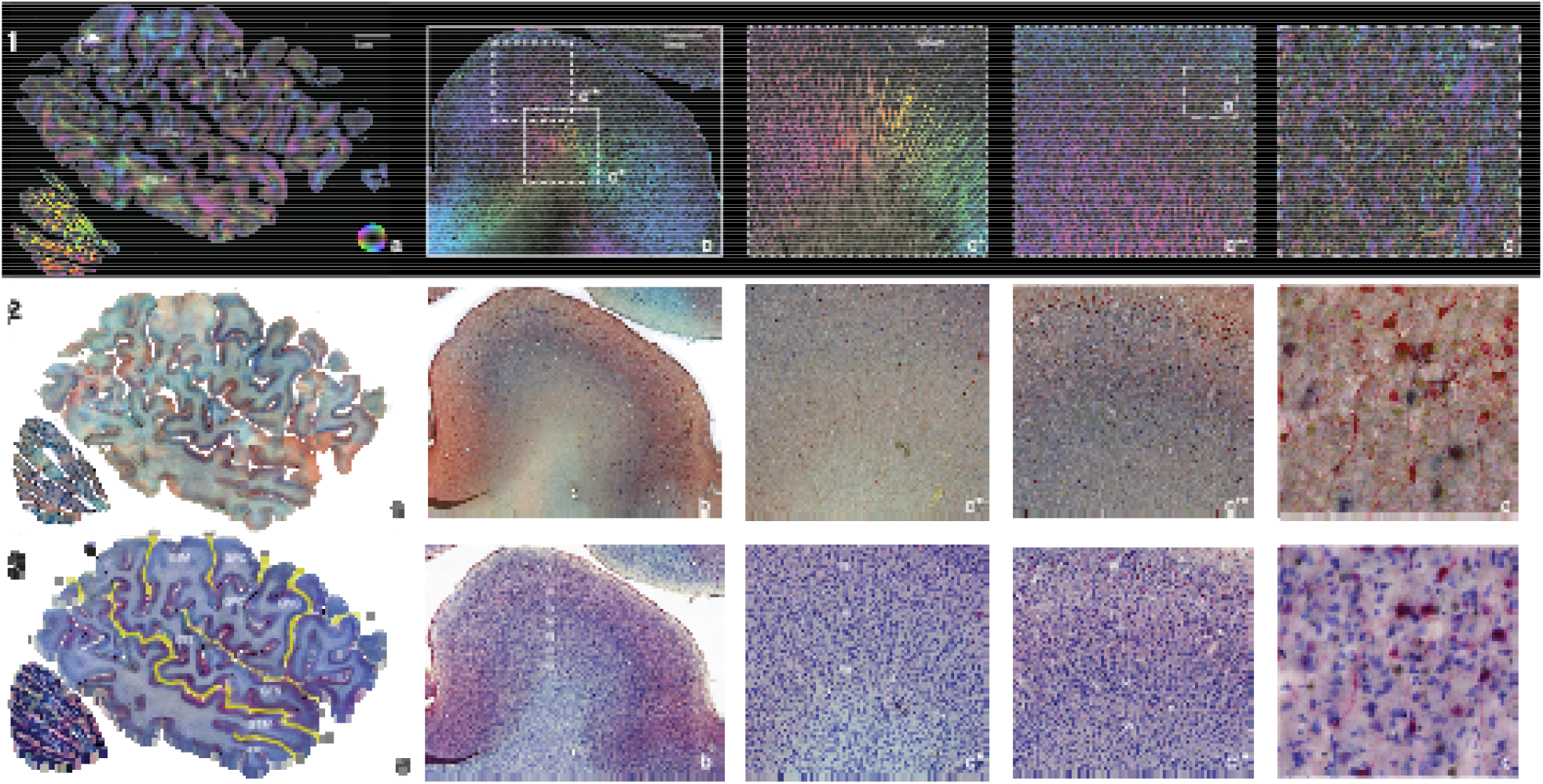
FACTs images of fibers (1), parvalbumin (blue), calbindin (magenta) and calretinin (yellow-brown)-expressing cells labeled using chromogenic immunohistochemistry (2), as well as Cresyl violet cytoarchitecture (3), in the same, full, sagittal human brain section (a). Detail panel (b) shows complementary layering patterns in area PG of the posterior inferior parietal lobule. Higher (same) magnification images for layers IV (panels c*), and III (panels c**) from panel b, for each corresponding modality. For the calcium-binding protein staining we present (2), the richest signal in cell body densities (2c**) is located at complementary locations to the richest fiber densities (1.c*). This effect is likely dependent on the targeted protein but serves to illustrate the supra-additive contribution of the multimodal signal acquired to the cytoarchitectural definition, of similar overall density (3.c* & 3.c**). Panels d showcase micron-resolution details of panels c** for each modality at the same location, with 2.d revealing sparse full cell morphologies for molecularly identified cell types (2.d), with labeling for all molecular targets (dotted circle) as well as distinct neural processes (arrows), complemented by Cresyl violet cytoarchitecture (3.d) and sparse myelinated fibers surrounding *-marked cell exclusions (1.d). The inclusion of full morphologies in the imaged data is a matter of probability, dependent on cell orientation, cutting plane, section thickness, and imaging focus. Labeled relevant landmarks are, clockwise, sulci: fpo – fissura parietalis occipitalis, si – sulcus intraparietalis, spc – sulcus postcentralis, sc – sulcus centralis, sprc – sulcus precentralis, sfi – sulcus frontalis inferior; fl – fissura lateralis, sts – sulcus temporalis superior, sti – sulcus temporalis inferior, gyri: GSM – Gyrus Supramarginalis GC – Gyrus Centralis, GPrC – Gyrus Precentralis, GTS – Gyrus Temporalis Superior, GTM – Gyrus Temporalis Medius, GTI – Gyrus Temporalis Inferior, GTT – Gyri Temporales Transversi, and cytoarchitectonic areas PG of the LPI – Lobulus Parietalis Inferior. Marked locations in panel 1.a indicate location of details magnified in Figs. 3–5. Scale bars for each magnification represent the size in the imaged section, rather than original brain space. Size bars are 1 cm for panels a, 1 mm for panels b, 500 µm for panels c*&c**, and 100 µm for panels d, respectively.

### Tissue preparation and sectioning

The method is applicable to formaldehyde-fixed tissue, whether fixed in formalin or in paraformaldehyde (PFA), making it accessible for regularly preserved human post-mortem tissue, as well as experimental animal tissue. For a full human brain, fixation requires 16 weeks submersion in 4% formalin, followed by washing in running tap water for 1 week to remove excess fixative. The tissue is subsequently moved to phosphate buffered saline (PBS; 0.1M, pH7.4) with 0.1% azide (NaN3). The meninges are removed, and the brain sectioned along the midline into two hemispheres, releasing air trapped in the tissue facilitate further tissue preparation and imaging. T2* weighted MR images (Philips Ingenia 3.0T CX) are acquired to correct for tissue distortions and to facilitate reconstruction of the full brain volume8, for digital atlas registration^31^.

Following intact brain imaging, the tissue is immersed in 20% glycerol in PBS (0.1M, pH7.4, + 0.1% NaN_3_), at room temperature. When sinks, it is frozen at -70°C as a whole (hemisphere), and sectioned (sagittal, 50 µm) on a Leica Biosystems CM3600 freezing sliding microtome. The section thickness of 50 µm enables safe handling of the full free-floating sections and also corresponds to the depth of field of a flat scan at 1 µm in-plane resolution imaging with bright field (see later section). Block-face images are acquired for each section before cutting and collecting on a glass microscopy slide to produce an undistorted reference image. The tissue (dry/frozen) is stored on glass slides at -80°C. Tissue stored this way remains viable for imaging and processing for an extended period (years) and allows us to progressively extract individual sections for further processing.

We tested parts of the protocol (Supplementary Table 1) in full hemisphere human sections cut in the frontal plane, and validated the protocol in single sections for both rodent (rat – Rattus norvegicus, Fig. 4). and in non-human primate (vervet - Chlorocebus sabaeus, Fig. 5) brains. The animal specimens were euthanized according to standard protocols (see Online Methods) and fixed by perfusion. All subsequent steps were identical to the protocol reported for human.

### 3D Polarized Light Imaging

The unstained sections are coverslipped in 20% glycerol in PBS for imaging. 3D Polarized Light Imaging (3D-PLI^32^) is used to image label-free the largest surfaces (up to 180 mm^2^) with an in-plane resolution of 1.3 µm × 1.3 µm. The measurement relies on a contrast mechanism that is intrinsic to the tissue, i.e. birefringence, an optical property of anisotropic materials such as brain tissue with its intricately interwoven axonal projections^32^. The basic principle of 3D-PLI is to pass polarized light through a thin brain section, and measure alterations in the polarization state of the light. This generates contrast between structures with different optic axis orientations. The optic axis of the tissue is determined by the orientation of axons. When the polarizing filters are being rotated, a signal is measured that has a sinusoidal shape. It is used to estimate the local orientation of the axons in each measuring field^11^. After 3D-PLI, the coverslip is removed in a PBS bath, and the sections are carefully unmounted, to continue to with the immunohistochemistry procedure.

### Immunohistochemistry for molecularly identified cell types and cytoarchitecture

We label parvalbumin (PV), calbindin (CB) and calretinin (CR) calcium-binding proteins in free-floating, full human brain sections (frontal, or sagittal cutting planes). To achieve high signal-to-noise ratio, as well as signal amplification, we use a primary, secondary, as well as a tertiary chromogenic development step for each targeted protein (see Online Methods). Optically distinct chromogens (yellow-brown diaminobenzidine for CR, the alkaline phosphatase (AP) substrate Red AP (Vector SK-5100), which yields a magenta reaction product, for CB, and Blue AP (Vector SK-5300) for PV) are used in order differentiate the three cell types. The sections are then cleared in a xylene-free system, to preserve AP signal.

High-resolution (1 µm in plane) brightfield images of the molecularly identified cell types are acquired using a Huron TissueScope™120 slide scanner. In the next step, the slides are immersed in xylene to remove the glass cover and destabilize AP chromogens staining for PV and CB-positive cells. Subsequently, the tissue is stained with Cresyl violet (Nissl), cleared with xylene, and re-coverslipped. The sample is re-scanned to add cytoarchitecture, in the same location (and same mount).

### Tissue deformation

Two geometric reference points for image registration are collected: one volumetric – structural MR, and one planar – cutting blockface. The former is acquired in PBS, after the tissue was first fixed and washed. The latter is acquired from the frozen tissue embedded in 20% glycerol in PBS solution. The tissue is subsequently mounted in the same glycerol solution. Since the tissue is floating, the section alignment in the PLI acquisition may vary from the blockface, but not the absolute surface. The section is then re-floated in PBS, immunohistochemically stained, manually re-mounted on gelatin coated slides using the blockface image as a reference, and air dried. As a result of this processing, the alignment of the section is changed from the PLI mounting, and the section surface is shrunk to 93% of the original size axially, and to 85% as a function of surface. A volumetric calculation is not fully applicable, as the tissue is sectioned, and the imaging plane of each section is a projection of the 50 µm section thickness. After the chromogenic immunohistochemical staining and drying, the specimen has been permanently altered and can no longer be removed or repositioned. All subsequent steps, that involve strong solvents and dehydration (xylene and ethanol) do not affect the surface of the section. The re-stained Cresyl violet image has identical alignment and size with the immunohistochemistry image.

Since the image acquisition is done in the same complete section for the different modalities, the shrinking loses its relevance for topographical comparisons and registration, as the images can be aligned in their entirety based on pervasive inherent features, such as sulcal pattern, and/or blood vessel exclusions. The images in Figs. 2–5 have been manually aligned in Adobe Illustrator 2026, for illustration purposes. Example images acquired with the FACTs method are available from the authors upon request (see Data availability); their size is, depending on the section, 10 Gigabyte to 40 Gigabyte.

### Identification of anatomical structures: areas and tracts

The same-section workflow allows direct evaluation of the widespread practice of integrating histological modalities or assigning areal borders in adjacent sections (typically separated by 20-50 µm). In a region of the intermediate postcentral sulcus where lamination is partially tangential to the cutting plane (Fig. 3), modality-specific borders of cortical areas align consistently within a given section but shift across consecutive sections. This demonstrates that features visible in one modality can be reproduced, but cannot be assigned to cytoarchitectonic delineations derived from neighboring sections with high spatial accuracy due to their shift.

**Fig. 3.**
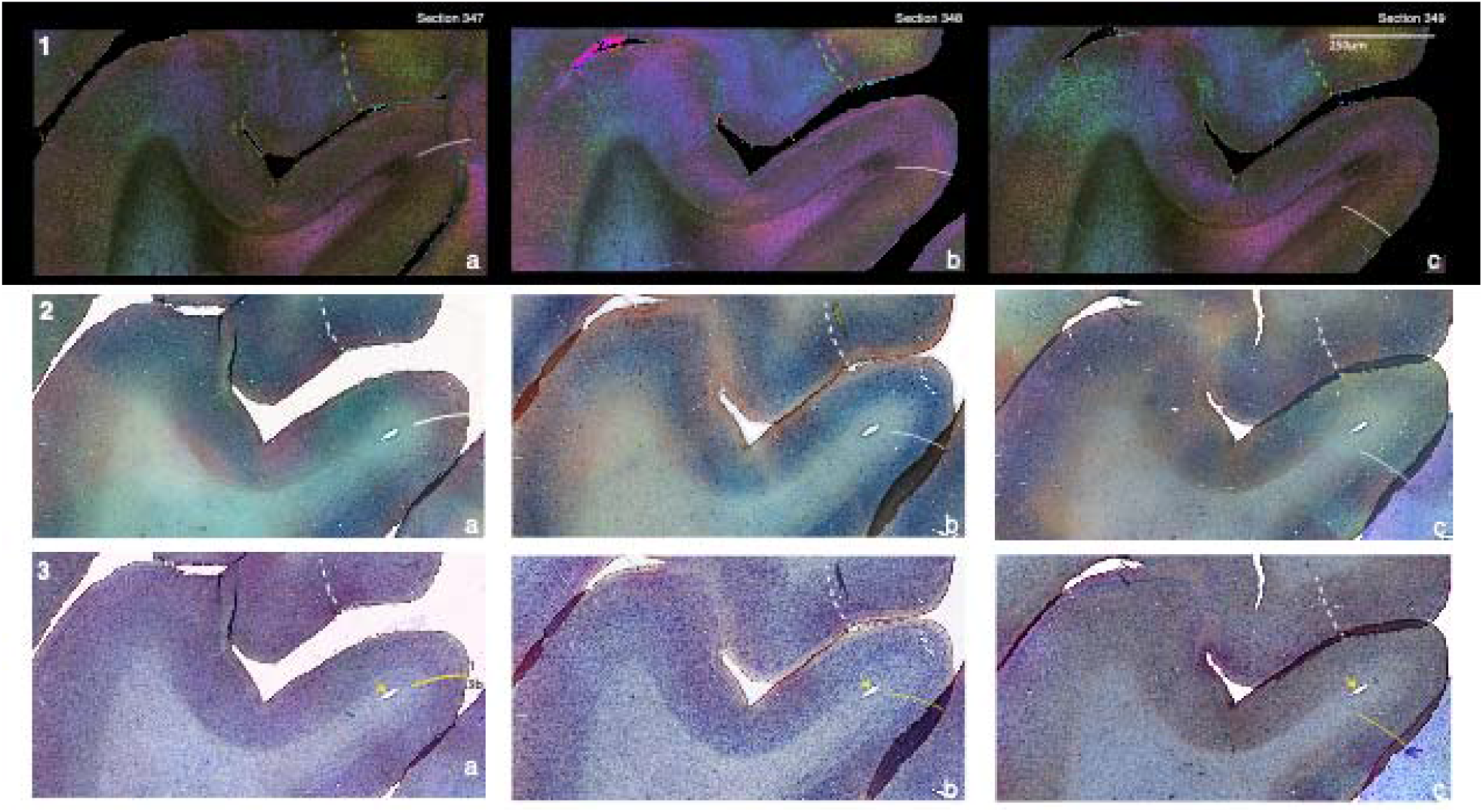
FACTs images of fibers (1), molecularly identified cell types (2) and cytoarchitecture (3) in the postcentral sulcus (location marked in Fig. 2.1.a), in three consecutive sagittal sections: the reference section 348 (panels b), the preceding, lateral section 347 (panels a) and the subsequent, medial section 349 (panels c). Solid yellow lines in panels 3 indicate the border between areas 3b and 1: area 3 is more cell dense and has smaller pyramidal cells in layer III^50^. Solid white lines in panels 1 and 2 show the same boundary superimposed on the fiber and cell-type modalities. Dashed lines (green in panels 1, white in panels 2 and 3) mark a transition that can be observed in the fiber modality, which also coincides with corresponding features in the cell-type and cytoarchitecture panels, when superimposed on these modalities. This transition as an example of the additional information afforded by multimodal same-section imaging. Laminar patterns are similar across the three sections in all modalities, attesting to the reproducibility of the workflow. At the same time, the architecture changes slightly over the 150 µm spanned by sections 347–349. Accordingly, the position and the angle of the area 3b/1 border relative to a blood vessel (yellow arrows, panels 3) change from section to section; between sections 347 and 349, the boundary is displaced by approximately 100–150 µm and rotated by approximately 35–45°, with section 348 occupying an intermediate position. Same magnification for all panels; the scale bar (250 µm) represents dimensions in the imaged section rather than in the original brain space.

The three modalities reveal partially overlapping and complementary relationships. In a full rat sagittal section (Fig. 4), compact parvalbumin-positive fibers of different sizes are embedded within the mammillothalamic tract. These observations provide information on the internal organization and relative size of chemically defined projections that is difficult to obtain with conventional tracing combined with immunohistochemistry.

### Similarities and differences of laminar pattern across the three modalities

Laminar patterns likewise show both correspondences and divergences across modalities. At the boundary between primary visual cortex and the optic tract in a vervet sagittal section, and within Heschl’s gyrus in the human, laminar borders exhibit alignment (visible in all three modalities), partial correspondence (shared by two modalities), or modality-unique boundaries (Fig. 5). Since classical cytoarchitectonic layers are defined by Nissl staining, borders observed in fiber architecture or molecular markers should be interpreted as related but not necessarily identical laminar features. The multi-modal view offered by FACTs therefore enables the mapping of more complex laminar and sublaminar patterns by integrating complementary information from different structural domains.

### Transferability across species

To assess its transferability, the FACTs workflow was applied to full sagittal sections of a Wistar rat and a vervet brain. In contrast to human post-mortem material, both animal brains were perfusion-fixed under terminal anesthesia and were therefore not subject to a post-mortem delay. Once fixed, the tissue was processed with a virtually identical protocol across species. The only difference concerned the duration of cryoprotection: owing to their smaller size, the animal brains equilibrated in the glycerol solution considerably faster. Because complete sinking of the tissue defines the endpoint of this step, the resulting preparation is equivalent irrespective of brain size. Labelling and imaging procedures were identical to those used for human tissue (Online Methods). In the rat, fiber orientation, PV-, CB- and CR-expressing cells and cytoarchitecture were acquired in the same section, from the whole-section overview to the level of individual myelinated bundles within the mammillothalamic tract (Fig. 4). In the vervet monkey, the three modalities were likewise obtained across the full section, resolving the optic radiation and the adjacent cortex at cellular resolution (Fig. 5). These examples demonstrate that FACTs provides an identical methodological basis on which comparisons across species can be conducted; they do not themselves constitute an interspecies comparison, which would require sampling of homologous regions in multiple individuals per species.

**Fig. 4.**
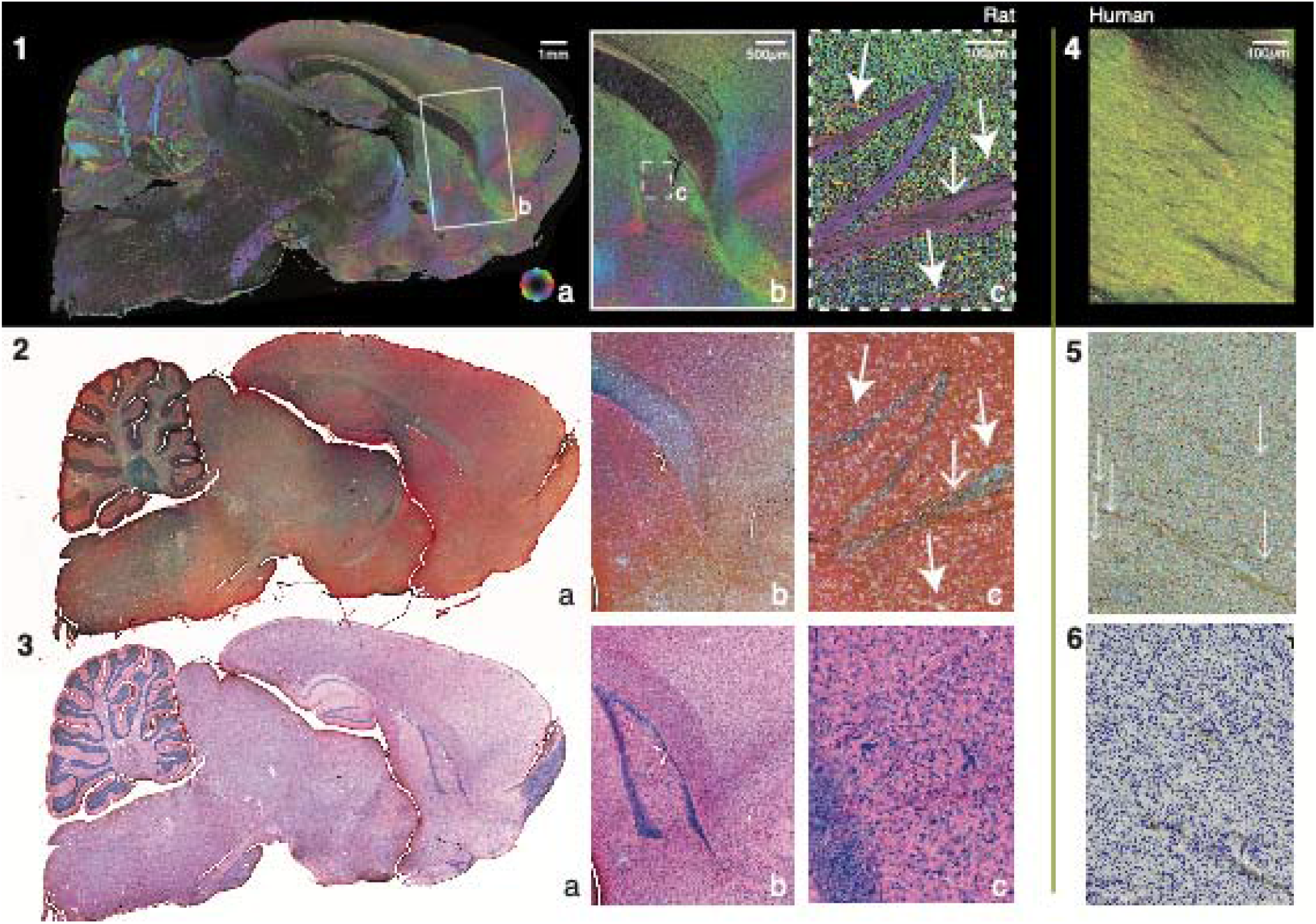
**FACTs validation in a Wistar Rat (Rattus** norvegicus) full brain sagittal section, revealing fiber orientations (1), parvalbumin (PV, blue), calbindin (CB, magenta) and calretinin (CR, yellow-brown)-expressing cells (2), as well as Cresyl violet-stained cytoarchitecture (3). Panel a of each modality shows full section overview, while panels 1.b through 3.b showcase a magnified view of the corpus callosum together with the fornix and mammillothalamic tract (MTT) in rat (1.b) complemented by corresponding calcium binding protein expression (2.b) and cytoarchitecture (3.b). Panels 1 through 3.c further zoom into the MTT, revealing myelinated bundles (1.c, solid arrows), including a subset of PV-positive projections (2.c, open arrow), in the context of local cytoarchitecture (3.c). Panels 4 through 6 showcase an example in human where CB-immunoreactive single axons (arrows) can be identified within the inferior longitudinal fasciculus (inset Fig. 2.1.a). Scale bars for each magnification represent the size in the imaged section, rather than original brain space. Size bars are 1 mm for panels a, 500 µm for panels b, 100 µm for panels c, and 100 µm for panels 4 through 6, respectively.

**Fig. 5.**
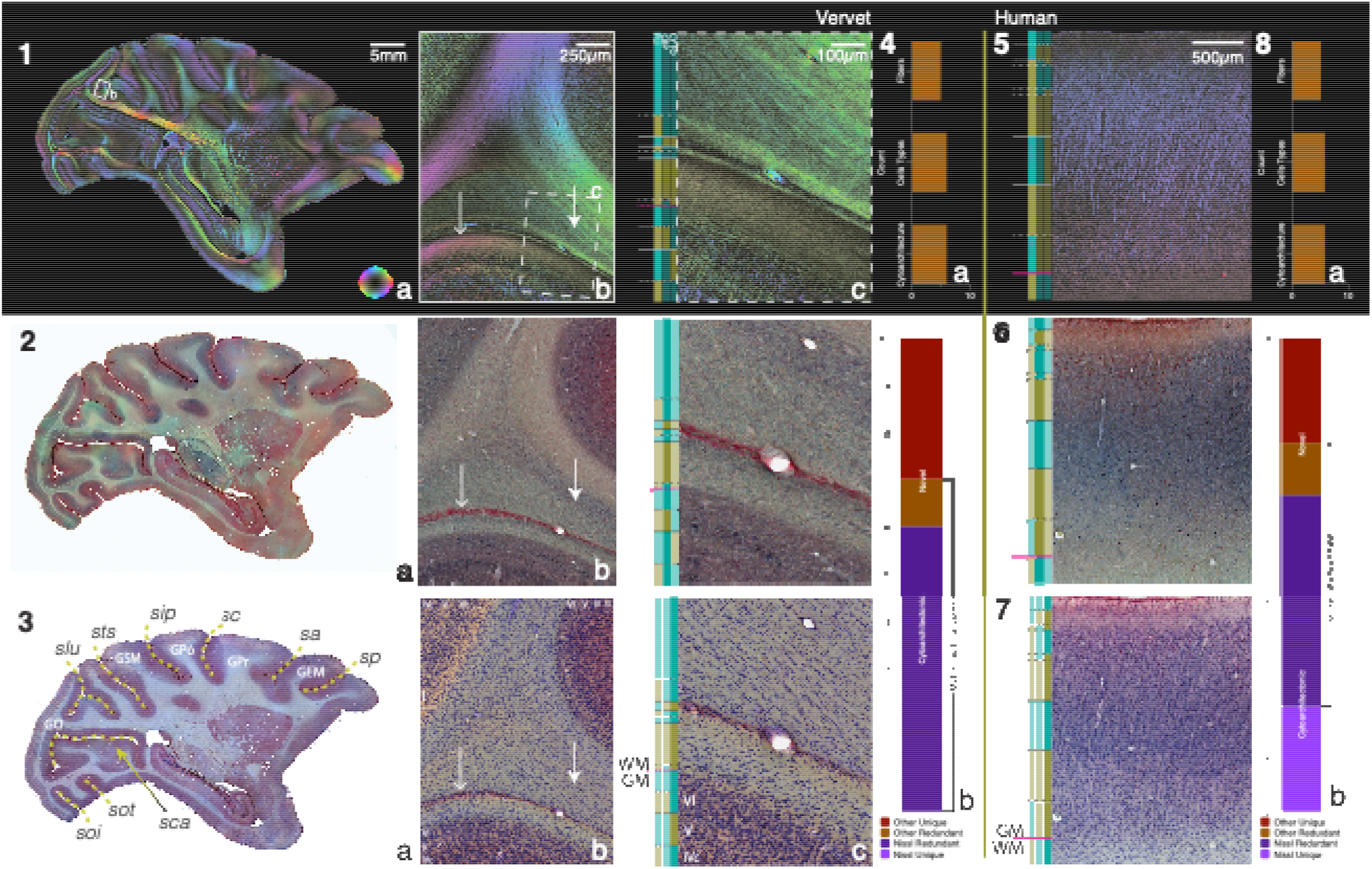
**FACTs validation** for Vervet (Chlorocebus sabaeus) showing fiber orientation map (1), parvalbumin (PV, blue), calbindin (CB, magenta) and calretinin (CR, yellow-brown)-expressing cells (2), as well as cytoarchitecture (3). In panel 3, the main landmarks indicated are sulci, from the posterior aspect, clockwise: sca – sulcus calcarinus, soi – sulcus occipitalis inferior, sot – sulcus occipitotemporalis, slu – sulcus lunatus, sts – sulcus temporalis superior, sip – sulcus intraparietalis, sc – sulcus centralis, sa – sulcus arcuatus, sp – sulcus principalis, and gyri: GO – Gyrus Occipitalis, GSM – Gyrus Supramarginalis, GPo – Gyrus Postcentralis, GPr – Gyrus Precentralis, GFM – Gyrus Frontalis Medius, GFI – Gyrus Frontalis Inferior. Panels 1.b through 3.b showcase a magnified view in the optic radiation at the position indicated in panel 1.a, revealing diverging fiber orientations (1.b), including a subset of PV-positive projections (closed arrows), as well as a narrow strip of CB-positive cells (open arrows). All FACTs features together are organized in distinguishable bands with a higher complexity than the aligned local cytoarchitecture (3.b). Panels 1.c–3.c show higher magnifications of the structures indicated in panel 1.b. Panels 5 through 7 are FACTs images corresponding to the location indicated in Fig. 2.1.a, showing human primary auditory cortex (Heschl’s gyri / GTT). Vertical color vectors along panels 1.c-3.c and 5-7 indicate visually identifiable changes in lamination for each of the imaged modalities, with yellow-teal switches. For each modality the corresponding vector is high intensity colored, while the corresponding other are low intensity. The vector order is preserved, as marked in panel 1.c. Horizontal solid lines denote lamination borders shared by all three imaged modalities, densely-dotted lines denote borders visible in two of the imaged modalities, while sparsely-dotted lines correspond to lamination borders only visible in one modality. The magenta horizontal line marks the white matter (WM) / gray matter (GM) boundary, as typically defined. Panels 4 and 8 summarize the identifiable borders for the illustrated vervet and human examples, respectively with (a) the number of switches visible in each modality, and (b) the correspondence between modality borders marked as redundant. Scale bars refer to dimensions in the imaged section: 5 mm (a), 250 µm (b), 100 µm (c and 5–7).

## Discussion

Maps of the human brain are valuable tools for gaining a deeper understanding of the biology behind brain function. Here, we present a new workflow that produces multiple maps coming from a single section, expanding the scope of existing multimodal mapping. This workflow encompasses three aspects of human brain architecture: cytoarchitecture, fiber architecture and triple immunohistochemical characterization of cellular molecular expression. Resulting full-section data reveal macro-scale correspondence between different modalities (Figs. 2, 4 and 5), with full cytoarchitectonic embedding, novel meso-scale features such as fully stained molecularly defined projections, together with the complete local white matter, and extended lamination information in the cortex, as well as cell morphologies and single axon connections. The complete topographical information, and micron-level cross-modal embedding remove the necessity of sampling or cross-specimen assumptions of correspondence, and open the possibility to explore multi-modal relationships in natively paired, and registered data.

The procedure has been tested for calcium-binding proteins parvalbumin, calbindin and calretinin. These three markers together account for the majority of cortical interneurons and have largely non-overlapping distribution patterns^34^. Calcium-binding proteins (CBPs) are a major class of proteins that differ in expression between (and can be used to classify) neurons^35^. These EF-hand proteins bind intracellular calcium, thereby directly influencing calcium signaling and all related neuronal functions, from ion channel opening to firing behavior to long-term potentiation. CBPs are expressed in all neurons, are essential for Ca2+ ion handling^36^, and thereby intrinsically related to the ionic balance and firing behavior of neurons, making their distribution relevant to their physiology and function. It can however be used with any set of up to three molecular targets. This limitation is inherited from the chromogenic, rather than fluorescent, immunohistochemical labeling chosen, which is on the other hand highly sensitive, and free of lifespan interference from the imaging procedure. It does not fade or bleach, in the absence of chemical agents, and can be coverslipped with a curing medium, which allows for long-term vertical storage at room temperature, i.e. it is space and energy effective. Additionally, multi-chromogen data can be acquired simultaneously for all immunolabeled targets, using brightfield scanning microscopy. Together, these ensure both uniform quality and speed, since no additional channel sequence is required. This is particularly relevant for large human brain sections. Of the 3 labels, two (Blue and Red AP) have high (di)methyl benzene sensitivity and can be broken in the tissue using xylene. The third label, DAB, is stable across treatments and can be used as an additional anchoring point between the two brightfield images (Fig. 1), cell-types and cytoarchitecture (Fig. 2). We opted for 3 chromogens that can be readily distinguished visually and separated in brightfield images, based on their different color/hue. All chromogens can be combined, and do not obscure the presence of another (unlike black DAB-nickel), allowing for simultaneous (faster) scanning, as well as the possibility to detect marker co-expression (double staining).

Two AP-bound and one HRP-bound secondary antibodies are used, followed by their specific chromogen development. Due to the shared substrate, the two AP-bound secondary antibodies need to be reacted and developed independently, to avoid cross-reactivity. To reduce the number of steps, the HRP-bound secondary antibody binding step can be performed together with either AP-bound antibody. Grouping the HRP and 1^st^ AP-bound secondary antibodies and developing them both before adding the second AP-bound antibody resulted in lower binding / unbinding of the primary antibody from the substrate, due to the repeated washes and temperature changes. The optimal configuration was to first react and develop one AP-bound secondary antibody, followed by amplification of the remaining primary antibodies and subsequent development of the other two secondary antibodies.

It must be emphasized that the entire process requires a several, precisely timed steps. The workflow makes use of numerous buffers (Online Methods, Supplementary Table 1), of varying ionic basis and pH, to allow for optimal chromogen development, as well as integration with 3D-PLI and Nissl staining. HRP and AP chromogens have specific pH requirements, which are intrinsically incompatible, with HRP development with DAB optimal between pH 7.2 and 7.5, with clear differences in color results, and extreme precipitation and background at higher pH, while AP development narrowly works at pH 8.4. The precise buffer composition, sequence, and pH are essential for ensuring stable and reproducible staining results, as the specific reactions of all substrates utilized are dependent on these. All development reactions are performed in the dark, as light exposure also influences reaction dynamics of all used chromogens.

The workflow is precisely balanced to maximally capitalize on both the strengths and vulnerabilities of the histological techniques integrated. As such, cresyl violet Nissl staining is obligatorily mounted, since the alcohol dehydration steps shrink and deform free-floating tissue, making it the natural last step of the procedure. At the other end is the 3D-PLI in untreated, label-free, mounted sections, making it an ideal first step, before adding any labeling agents to the tissue. Together with order, interaction reversibility of chemical agents employed in the procedure plays a major role in enabling the integration of histological techniques we use. The irreversible interactions such as the use of the dimethyl sulfoxide (DMSO) in earlier 3D-PLI protocols^11^ have been removed because it inhibits alkaline phosphatase oxidation. It has been replaced with room-temperature 20% glycerol in PBS incubation, with 0.1% NaN3. This switch has the additional advantage of substantially reducing incubation time from weeks, sometimes months^38^ to a couple of days. Adding NaN_3_ prevents adverse effects of room temperature incubation by inhibiting peroxidase activity and can be easily washed out of the tissue to prevent it from interfering with DAB staining.

By implementing these strategies, the presented workflow enables systematic and integrated mapping of protein expression in the human brain, complete with full topography and morphology. The immunohistochemical staining presented here can be expanded to any primary antibody set of choice, up to three simultaneous targets, with application ranging from complementary-protein-to disease-specific-maps. Since PLI and cytoarchitecture data acquisition are not specific for a certain cell-type, and not affected by changing molecular markers, the data are intrinsically anchored to cytoarchitecture by the Nissl stain, and can be easily compared to other datasets from other research groups based on this common denominator, as well as integrated into digital atlases^39^ and related to identified brain areas.

Multimodal, single-cell resolution, high-coverage human brain data is also available owing to the development of methods such as patch-seq^40^. This innovation bridges the gap between several modalities, such as transcriptomic data, together with electrophysiological recordings of single cells. Extensive data has been collated this way in resected blocks^41^, but spatial information, and cytoarchitectonic areal designation can only be semantic. In fact, any existing large surface/volume maps that integrate multiple modalities achieve only indirect topographical mapping, either in the same brain, by assigning different (alternating) sections to different data acquisition modalities^42^, or different brains^43^, by registering data sets acquired in different modalities^44^. These approaches rely on the premise that spatial correspondence between various features is robust, and replace direct spatial mapping with the assumption that either differences between sections are small enough to allow for inferring relations from data collated across rather than within section, or that cross-brain registration is sufficiently accurate at a micron level. Such assumptions have not been directly demonstrated, leaving spatial relationships between different architectural features unexplored.

Conversely, directly anchored multimodal images of (full) brain architecture can help disambiguate the boundaries and 3-dimensional configurations of architectural features, by providing additional/complementary structural insight in complex configurations of multiple converging boundaries and cutting planes non-orthogonal to the structure of interest^45^. Fig. 3 illustrates incremental shifts in boundary location between consecutive 50 µm-thick sections and challenges the assumption of micron-level correspondence between adjacent sections. The region shown is anatomically complex: at the intermediate postcentral sulcus, the cortical surface of BA1 bends sharply from the transverse into the sagittal plane, so that the cutting plane intersects the cortex at a rapidly changing angle. This renders the identification of areal boundaries from cytoarchitecture alone particularly difficult, and complementary information from additional modalities acquired in the same section may help to resolve them. Although Fig. 3 depicts a region in which lamination runs partially tangential to the cutting plane, this configuration is not exceptional: in the highly folded human cortex, oblique and tangential cuts occur in every whole-brain section, irrespective of sectioning orientation. Combined with section-specific non-linear deformation, and with a spacing of 50 µm that exceeds the diameter of most neuronal somata, this renders cross-section correspondence an approximation at the micron level in any brain region, whereas same-section acquisition removes this source of error altogether.

The method we present is reliable for full (sagittal) human brain sections, it is robust and reproducible in several human samples as well as typical animal-model brains (both smaller and fixed under higher-control conditions than human donor tissue), without individual adjustments. The features we image are in line with previous findings / maps, where these exist. We go beyond existing maps by achieving two major advancements in brain mapping: micron-resolution full (section) mapping, coupled with true multimodal acquisition, in the same tissue, for different modalities, bridging modalities and scales. It was necessary to modify the techniques for individual components to make them fit different modalities together with veritable topographical anchoring. On a middle term this workflow allows to generate large numbers of numerical, multimodal, high-resolution maps of cortical architecture, by joining connectivity, cell type identification by (triple-labeled) protein expression, and cytoarchitectural anchoring, at the whole human brain level. The potential to expand human brain maps with the addition of morphologically complete (non-sampled) molecular information fully topographically embedded with both cytoarchitecture and connectivity can open new opportunities for understanding brain organization with impact ranging from medical research application to modelling, using supercomputing and big data analysis^46^.

## Online Methods

### Tissue preparation

For the full brain human experiments, we acquired intact donor brains and hemispheres (Ethical approval 2017-1037 NBB) from the Netherlands Brain Bank, from psychiatric and neurological controls. The postmortem delay for all specimens was shorter than 6 hours (Supplementary Table 1). After the autopsy, we fixed the tissue by immersion in 4% formalin for 16 weeks. This relatively short fixation period ensures the tissue can be successfully processed for immunohistochemistry. We consistently remove the tissue from the fixative at a set time point, to ensure uniform quality across specimens.

For the animal model validation, we used tissue from an adult Wistar rat that was perfusion-fixed with 4% PFA under isoflurane anaesthesia and subsequently decapitated. Animal tissue procedures and husbandry were approved by the institutional animal welfare committee at the Research Centre Jülich and were carried out in accordance with the guidelines of the European Communities Council Directive for the care and use of animals for scientific purposes. Tissue was also used from an adult male vervet monkey from the Wake Forest University (Winston-Salem, NC, USA) that was perfusion-fixed in accordance with the UCLA Chancellor’s Animal Research Committee (ARC) ARC #2011-135 and by the Wake Forest Institutional Animal Care and Use Committee IACUC #A11-219. After perfusion, the brains were carefully removed from the skull, cryoprotected by immersion in 20% Glycerol in PBS with 0.1%NaN_3_, at room temperature, until sinking, deep frozen in isopentane at[]−50°C, and placed in pre-cooled plastic bags and stored in a deep freezer at − 80 °C until further sectioning.

### 3D-PLI preparation

Unstained whole-human brain sections (tissue area: 10 cm × 15 cm) were scanned using polarizing microscope LMP-1 (Taorad GmbH, Aachen, Germany). The device has a field of view of 2.7 mm × 2.7 mm and provides large-area tile-wise scanning with a pixel size of 1.3 µm to derive the local orientations of myelinated axons at the microscale. The intensity of the light transmitted through the brain section placed between a rotating linear polarizer and a circular polarizer follows a sinusoidal course, is referred to as light intensity profile. Therefore, scanning is done at discrete equidistant filter rotation angles of 20° generating stacks of nine images per tile. In sum, the polarization measurement of one entire sagittal human brain section (Fig. 2) comprises about 22,000 images leading to 0.8 TB of raw data (per section). The scan time of such a section is currently 12 hours.

An HPC-based analysis pipeline allows to derive complementary parameter maps^47^ from the light intensity profiles by means of the Jones calculus for linear optics^48^ and discrete Fourier analysis^49^. The first, transmittance map is a voxel-wise measure for the extinction (scattering and absorption) of light after passing through the brain tissue. It provides images reflecting particularly the densely packed myelin. The second, retardation map, represents the normalized amplitudes of the light intensity profiles and reflects both, the birefringent strength as well as the out-of-sectioning-plane (z) orientation of the fibers. Thirdly, the fiber direction (φ) map describes the within-sectioning-plane (x/y) orientation of each fiber. And finally, the fiber inclination (α) map describes the out-of-sectioning-plane angle of each fiber.

The fiber direction and inclination maps are combined in the fiber orientation maps (FOMs), color-coded in the Hue–Saturation–Value (HSV) space, a cylindrical-coordinate representation (hexcone) of the Red-Green-Blue space. The HSV describes hue, saturation, and value as the three dimensions of a color. Hue is the dimension normally called red, yellow, blue, etc. Saturation specifies the departure of the hue from white or gray, and value indicates the departure of a hue from black. In this study, the focus was on the fiber sensitive FOM (Figs. 2–5).

### Immunohistochemistry

After 3D-PLI acquisition, we re-floated the sections by removing the coverslip seal for each slide, while submerged flat in a large PBS bath. All tissue pieces were then collected using a soft brush, and stored in PBS with 0.1% NaN3. Each full section was then individually stained free-floating, using chromogenic immunohistochemistry targeting calcium-binding proteins parvalbumin (PV), calbindin (CB) and calretinin (CR).

We first washed the sections (2x 5min) with PBS. We then blocked endogenous peroxidase activity with a 6% H_2_O_2_ (30%) in PBS solution (30 minutes). Following the blocking step, we removed any unreacted traces of peroxide from the tissue with repeated PBS washes (5 x 10 minutes). We blocked non-specific reactivity to the secondary antibody host using 2% normal donkey serum (NDS, 017-000-121 Jackson Immunoresearch) in PBS (+2% Triton™X-100 + 0.1% NaN_3_) for 1h. Subsequently, we incubated the tissue in the primary antibodies (SWANT PV 235, 1:1000; SWANT CB38, 1:1000; and SWANT CG1, 1:1000) in PBS (+5%NDS +2%TritonX +0.1%NaN_3_), first 10 minutes at 36°C, followed by overnight at room temperature, on an orbital tissue shaker.

The following day we switched the buffer system to Trizma®. After a 5 minute wash in Trizma® buffer (pH 7.4, 0.1M + 0.9% NaCl) we incubated the sections with the first secondary antibody (1:100 Alkaline-Phosphatase Donkey anti Mouse, 715-055-150 Jackson Immunoresearch) in the same buffer, for 1h at 36°C. We again washed the section for 5minutes in Trizma® buffer (7.4), followed by 5minutes Trizma® (pH 8.4, 0.1M + 0.9% NaCl) and developed it with the Blue AP (Vector SK-5300) chromogen kit (components 1/2/3 - 1/1/0.5:100 in Trizma® buffer pH 8.4 + 0.9%NaCl, 3 minutes, dark). We washed the section again (1 x 5minute in Trizma 7.4) and incubated it with the remaining secondary antibodies (1:100 Alkaline-Phosphatase Donkey anti Rabbit, 711-055-152 Jackson Immunoresearch, and 1:100 Peroxidase Donkey anti Goat, 705-035-147 Jackson Immunoresearch) in Trizma (pH 7.4 + 0.9% NaCl) for 1h at 36°C.

Following the second round of secondary antibodies, we washed the sample for 5 minutes with Trizma (pH 7.2 +0.9%NaCl) and developed the peroxidase with 3,3’ Diaminobenzidine tetrahydrochloride (DAB, D5905, Sigma-Aldrich, 10mg:20ml Trizma® buffer pH 7.2 + 0.9% NaCl, 30 seconds, dark), washed for 5 minutes in Trizma® buffer (pH 7.2 +0.9%NaCl), followed by another 5 minutes in Trizma® buffer (pH 8.4, + 0.9% NaCl). We then developed the remaining tag with the Red AP (Vector SK-5100) chromogen kit (components 1/2/3 - 1/1/1:100 in Trizma® buffer pH 8.4 + 0.9% NaCl, 5 minutes, dark), followed by a final wash with Trizma (0.1M, pH 7.4 + 0.9% NaCl).

### Mounting and brightfield imaging

We re-mounted the full brain sections on glass slides coated with gelatin (6% w/v + 0.1% w/v CrK(SO_4_)_3_ x 12H_2_O + 0.1% w/v NaN_3_), using Trizma® buffer (0.1M, pH 7.4 + 0.9% w/v NaCl). We used the blockface of the section (Fig. 1) as reference, to reassemble the floating fragments. Mounting of one section takes between 30 min and 4 hours, depending on size, and the number of fragments. We air dried the sections, then moved them to a 100% ethanol bath (20 minutes), followed by ClearRite3™ (20 minutes), and coverslipped with Leica CV Ultra (14070937891, Leica Biosystems). Tissue processed and mounted in this manner can be stored at room temperature for an extensive period (years) without color loss. After the Nissl staining (see procedure), we re-cleared the sections in xylene (2 x 10 minutes), and later re-coverslipped with Leica CV Ultra. While developing the method, we also coverslipped with VectaMount (H-5000) or VectaMount Express (H-5700-60, Vector Laboratories) for the chromogenic immunohistochemistry. We imaged the sections on a Huron TissueScope™120 Slide Scanner, with 10x magnification and brightfield illumination, generating 1 µm resolution full-section images. Both the cell types and the cytoarchitecture (Fig. 2) for a mid-sized section (section 348) required 4h for a single plane at 1 µm resolution, amounting to an average of 30GB of data per plane for each (CBP and cytoarchitecture) labels. To go towards true 3-dimensional micron resolution, we acquire 59 planes in each (50 µm) section, for both immunolabeling, and Nissl staining, for later image deconvolution.

### Nissl staining

We capitalized on the AP sensitivity to xylene, to (partially) remove the AP signal, and add further cytoarchitectural information with Cresyl violet Nissl staining. We submerged the slides in xylene for 1 week, which causes the glass cover to detach, and destabilizes the AP chromogen binding making it possible to label cytoarchitecture, without loss of signal from over-dense labeling (Fig. 3). The removal of the chromogens is partial: it is most pronounced for the Blue AP product, less pronounced for the magenta Red AP product, and absent for the DAB product, which instead darkens from yellow-brown to brown over the subsequent pH changes (Fig. 2). The resulting reduction in optical density ensures clear visibility of the Cresyl violet staining (Fig. 2, panel 3.d). Subsequently, we rehydrated the sections by immersion in a reverse alcohol series – 10 minute 100%, 70%, 50% ethanol, followed by 10 minutes deionized water, stained for 20 minutes in Cresyl violet 0.025%, differentiated with 50% ethanol + 0.025% v/v glacial acetic acid in a moving bath – 10 minutes, re-dehydrated with an increasing ethanol series – 10 minutes 70%, 100%, and finally 2x 10 minutes xylene (different baths), before coverslipping with Leica CV Ultra (14070937891, Leica Biosystems), earlier with DPX (1005790500, Sigma-Aldrich).

## Data availability

The original full-resolution images acquired with the FACTs workflow (3D-PLI, multiplex chromogenic immunohistochemistry and Cresyl violet Nissl staining) are stored at Forschungszentrum Jülich but currently cannot not publicly deposited, as their file size makes them impractical to transfer, load and view. They are available from the corresponding authors upon reasonable request and can be provided at a size and resolution specified by the requester.

## Acknowledgements

We thank Markus Cremer, Anna Steffens and Philipp Schlömmer for their technical assistance with histology and tissue imaging, and Felix Matuschke for his support with image processing.

