## Supplemental materials for "Brain FACTs: A same-section workflow for Fiber Architecture, Cytoarchitecture and cell Types in large-scale brain sections" (from another version of this preprint: version 1, 2026-10-15)

**Computational Workflow**

We have developed an efficient computational workflow to facilitate the management and HPC-based analysis of multimodal, petabyte-scale whole-brain datasets.

This workflow has been optimised for use with JURECA-DC (Jülich Research on Exascale Cluster Architecture), a modular pre-exascale supercomputer operated by the Jülich Supercomputing Centre at Forschungszentrum Jülich. The system consists of a flexible data-centric (DC) module based on the Atos BullSequana XH2000.
(https://apps.fz-juelich.de/jsc/hps/jureca/configuration.html)

The workflow comprises five main components:
1. Data acquisition and transfer: TIFF and BigTIFF images are moved from different microscopes (polarisation and bright-field) via laboratory PCs to a local buffer system and finally to the Jülich Supercomputing Centre.
2. Data conversion and quality control: Raw images are consolidated into HDF5 files and checked for acquisition errors.
3. Standardised data organisation and ingestion, including metadata validation, multiresolution image pyramids, automated insertion into the central data structure, and database indexing;
4. HPC-based analysis and reconstruction: SLURM-orchestrated processing chains (https://slurm.schedmd.com/overview.html) perform imaging-specific tasks such as colour normalisation, calibration, signal analysis, stitching, inclination estimation, segmentation, and fibre orientation mapping.
5. Data management and access comprises archival backup, workflow monitoring via a dashboard, metadata-driven searches and web-based visualisation.

**Supplementary Tables**

| Brain | Sex | Age | PMD | Section coverage | Sectioning plane | Modalities |
|---|---|---|---|---|---|---|
| Human 1 | F | 94y | 4:50 | Right hemisphere | Frontal | Cell types / Nissl |
| Human 2 | F | 76y | 5:30 | Left Hemisphere | Frontal | Cell types / Nissl |
| Human 3 | F | 68y | 3:30 | Left Hemisphere | Frontal | Cell types / Nissl |
| Human 4 | M | 86y | 4:15 | Whole brain | Sagittal | Fibers / cell types / Nissl |
| Vervet | M |  | 0 | Whole brain | Sagittal | Fibers / cell types / Nissl |
| Wistar Rat | M |  | 0 | Whole brain | Sagittal | Fibers / cell types / Nissl |

**Supplementary Table 1:** Overview of basic information of specimens we tested including species: Human (homo sapiens), Vervet (Chlorocebus pygerythrus) and Wistar Rat (Ratus norvegicus), sex: female (F) or male (M), age, post-mortem delay (PMD, time elapsed between confirmed death and tissue collection/fixation), extent of each section surface, the sectioning plane (orientation of acquired section in brain space), and the tested modalities within section.

| **Buffer** | **pH** | **Molar concentration** | **Additions (W/V)** | **Use** |
|---|---|---|---|---|
| PBS | 7.4 | 0.1M |  | - Washes |
| PBS | 7.4 | 0.1M | + Triton | - H2O2 incubation for reacting endogenous peroxidase |
| PBS | 7.4 |  | + Triton<br>+ NaN3 | - Blocking serum<br>- Primary antibody incubation |
| Trizma | 7.2 | 0.1M | + 0.9% NaCl | - Incubation with DAB Tetrachloride (+ H2O2)<br>- Wash before and after DAB Tetrachloride incubation |
| Trizma | 7.4 | 0.1M | + 0.9% NaCl | - Wash after primary antibody incubation<br>- Secondary antibody incubation<br>- Wash after AP Substrate incubation |
| Trizma | 7.4 |  |  | - Final wash before mounting |
| Trizma | 8.4 | 0.1M | + 0.9% NaCl | - Incubation with AP Substrate kits<br>- Wash before incubation with AP Substrate kits |

**Supplementary Table 2:** Overview of buffers necessary for the chromogenic immunohistochemistry for molecularly identified cell types. Each has been carefully adapted for the precise sequence described in the Online Methods, to maximize efficiency of the chromogen binding, separate pH and substrate dependent reactions, and preserve tissue quality.
